# FBH1 and RAD54L directly interact and cooperate to drive replication fork reversal

**DOI:** 10.64898/2026.05.22.727240

**Authors:** Mollie E. Uhrig, Courtney N. Johnson, Jordi Gomez, Alexander V. Mazin, Claudia Wiese

**Author notes:** Graduate Program in Cell and Molecular Biology, Colorado State University, Fort Collins, CO, 80523, USA.

## Abstract

Replication fork reversal alleviates DNA replication stress and maintains genome stability. We previously showed that FBH1 and RAD54L cooperate to promote fork reversal in human cells, and that FBH1-dependent fork reversal requires the branch migration activity of RAD54L. However, the molecular basis of this cooperation remained unclear.

Here, we identify both a physical and functional interaction between FBH1 and RAD54L. We demonstrate that purified FBH1 and RAD54L interact directly and form a complex at stalled replication forks in cells. Mapping studies revealed that RAD54L Lobe 1 is critical for interaction with the FBH1 2B subdomain. In cells, FBH1-RAD54L complex formation is enhanced in the absence of RAD51. Consistently, purified RAD54L displays a stronger affinity for RAD51 than for FBH1.

Using biochemical reconstitution assays, we further show that FBH1 and RAD54L promote fork reversal more efficiently together than either protein alone, with maximal reversal observed when FBH1 acts before RAD54L. Collectively, our findings establish RAD54L as an essential functional partner of FBH1 in replication fork reversal and provide mechanistic insight into the sequential coordination of their activities.

## INTRODUCTION

Accurate and precise DNA replication is essential for maintaining genome integrity during each cell division. However, replisome progression is hindered by obstacles such as DNA lesions, DNA secondary structures, limited nucleotide pools, and transcription-replication conflicts, all of which can slow or stall replication forks (RFs) [1, 2]. Stalled RFs are vulnerable to resection, double-strand break (DSB) formation, and chromosomal rearrangements [3]. To alleviate stress associated with prolonged RF stalling, cells employ several pathways to restore RF progression including repriming, template switching, translesion synthesis, and RF reversal [4, 5]. RF reversal involves the annealing of the nascent strands and reannealing of the parental strands, thereby remodeling a three-way DNA junction into a four-way structure resembling a Holliday junction [6-8]. RF reversal, when regulated properly, maintains genome stability by repositioning DNA lesions back into duplex DNA for excision repair, prompting template switching, and potentially promoting recombination mediated repair and replication restart following cleavage of the reversed RF by substrate specific nucleases [4]. In contrast, unregulated RF reversal can lead to untimely RF degradation and genome instability [8-10].

Efficient RF reversal is directed by the RAD51 recombinase and the ATP-dependent DNA motor proteins SMARCAL1, ZRANB3, HLTF, and the DNA helicase FBH1 [9, 11-14]. In addition, we and others have shown that RAD54L, a dsDNA translocase in the SWI2/SNF2 family [15] with established functions in homologous recombination DNA repair (HR) [16-19], also mediates RF reversal in human cells [20, 21]. We showed that RAD54L contributes to RF reversal in both described RAD51-mediated RF reversal pathways [9, 20]. Interestingly, RAD54L engagement in RF reversal differs in each pathway. During RF reversal by SMARCAL1, ZRANB3, and HLTF, RAD54L functions independently of its Holliday junction branch migration (BM) activity. In contrast, RAD54L BM activity is essential for RF reversal in the FBH1 pathway. However, the mechanistic basis underlying the cooperation between FBH1 and RAD54L during RF reversal has remained unclear.

FBH1 is a 3′−5′ F-box DNA helicase [22] that catalyzes RF reversal *in vitro* and in human cells [23, 24]. In addition, FBH1 is an SCF (SKP-CUL1-F-box) E3 ubiquitin ligase that ubiquitylates RAD51, thereby limiting RAD51 association with chromatin [25, 26] and, thus, is predicted to restrain HR during replication stress [25, 27, 28]. FBH1-mediated ubiquitylation of RAD51 may also limit RF reversal and therefore requires RAD51-stabilizing proteins such as BRCA2 to counteract FBH1 [9]. Previous studies demonstrated that the E3 ligase function of FBH1 is dispensable for RF reversal, whereas the FBH1 helicase activity is essential [9, 23]. Current biophysical and biochemical models support a mechanism in which FBH1 unwinds the lagging strand at stalled RFs [24, 29]. Moreover, SCFFBH1 has been proposed to remain bound behind the RF junction and use its translocase activity to reanneal parental duplex DNA by continuously feeding the lagging strand template into the junction [24].

Despite recent advances in our understanding of FBH1 in RF reversal, the contribution of RAD54L to the FBH1 pathway remains incompletely resolved. Here, we show that the purified RAD54L and FBH1 proteins directly interact and are part of one protein complex at stalled RFs in human cells. Guided by AlphaFold 3 structural predictions and analyses of RAD54L domains expressed in human cells, we mapped the FBH1 interaction site to RAD54L-Lobe 1 (residues 304-354). In FBH1, we then mapped the RAD54L interaction site to residues I910, E911, and D912 of isoform 1. Using functional reconstitution assays, we show that RAD54L and FBH1 together promote RF reversal more efficiently than either protein alone. Moreover, order-of-addition experiments reveal that reversal activity is strongest when FBH1 is added to model RF substrates first, followed by RAD54L. Collectively, these findings significantly advance our understanding of the cooperative relationship between FBH1 and RAD54L in RF reversal and provide strong evidence that RAD54L is required for efficient execution of the FBH1-mediated RF reversal pathway in cells.

## RESULTS

### RAD54L selectively defective in BM activity rescues nascent strand degradation in cells with 53BP1 knockdown

The two RAD51-mediated RF reversal pathways are characterized by distinct RF protection factors [9]. BRCA2, FANCD2, and ABRO1 protect reversed RFs generated by SMARCAL1, ZRANB3, and HLTF, whereas 53BP1 protects RFs remodeled by FBH1 [9]. To further assess the impact of RAD54L’s BM activity on the FBH1 pathway, we analyzed nascent strand degradation in HeLa RAD54L knockout (KO), RAD54L KO+RAD54L, and RAD54L KO+RAD54L-BM cells, described previously [20, 30, 31]. Cells with RAD54L-BM express mutant RAD54L (*i*.*e*., RAD54L-4A/S49E) selectively defective in BM activity [32]. However, the RAD54L-BM protein retains the ability to stabilize RAD51 filaments, translocate on double-stranded (ds)DNA, and stimulate RAD51-mediated strand invasion [32]. Cells were transfected with 53BP1 or a negative control siRNA (Fig. S1A), subjected to consecutive pulse labeling with two thymidine analogs, and DNA fiber assays were performed after treatment of cells with 4 mM hydroxyurea (HU) for 5 h, which stalls essentially all RFs ([33]; Fig. 1A). Consistent with nascent strand degradation, IdU/CldU ratios were significantly lower in 53BP1-depleted RAD54L KO cells expressing wild type RAD54L (RAD54L KO+RAD54L) than in control-depleted cells (Figs. 1B, S1B). In contrast, no significant difference in IdU/CldU ratios was observed between 53BP1-depleted and control-depleted RAD54L KO or RAD54L KO+RAD54L-BM cells (Figs. 1B, S1B). These results are consistent with the expected defect in RF reversal in the absence of RAD54L [20]. These results further show that BM-defective RAD54L cannot catalyze RF reversal and, as such, protects nascent strands from degradation in the absence of 53BP1. As expected, expression of wild type RAD54L restored RF reversal proficiency, resulting in nascent strand degradation of reversed RFs that are not protected by 53BP1.

**Figure 1.**
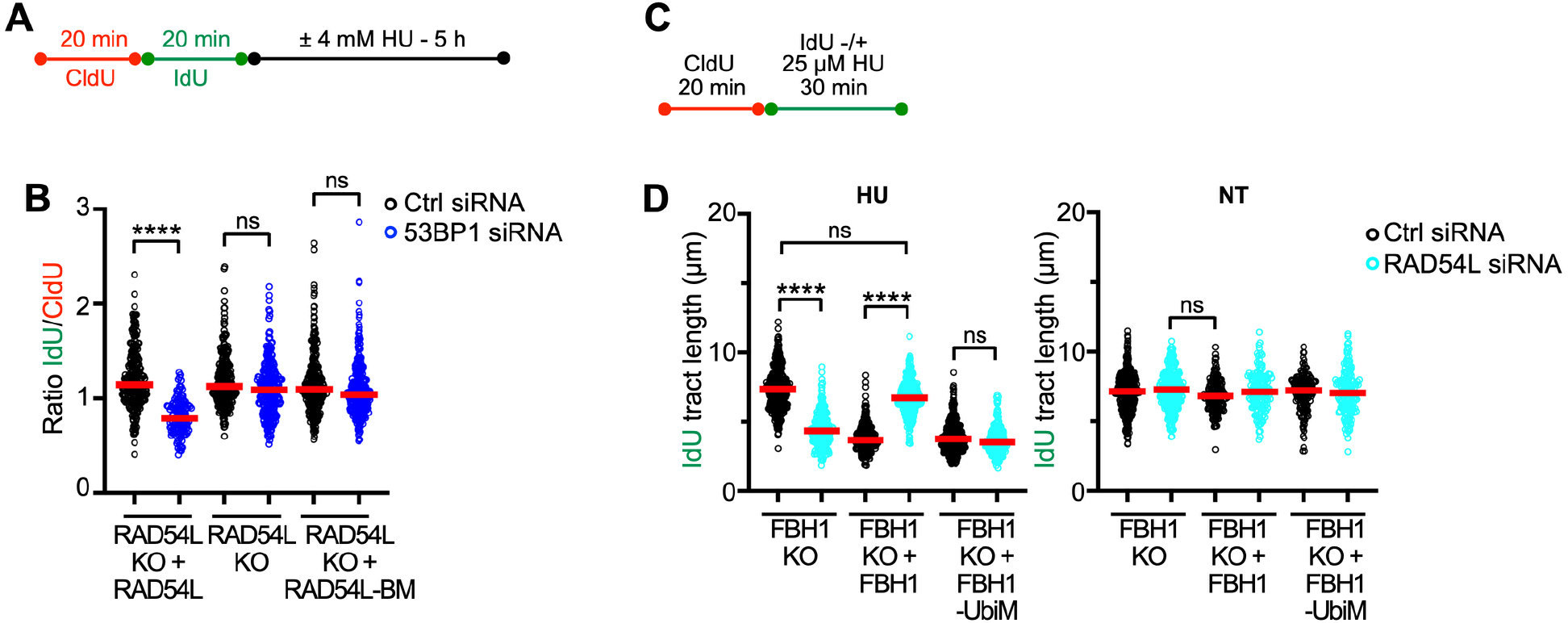
RAD54L branch migration (BM) is essential in the FBH1 pathway but dispensable for fork reversal by the FBH1 E3 ligase mutant. (**A**) Schematic of the DNA fiber assay protocol used in (B). (**B**) Dot plot with medians of IdU/CldU ratios in HeLa RAD54L KO cells and RAD54L KO cells with wild type RAD54L or mutant RAD54L-BM (*i*.*e*., RAD54L-4A/S49E [32]) and transfected with negative control (Ctrl) or 53BP1 siRNA (n=2; 124-318 fiber tracts analyzed; ns=not significant, ****p < 0.0001). (**C**) Schematic of the DNA fiber assay protocol used in (D). (**D**) Dot plots with medians of IdU tract lengths in HU-treated and untreated U2OS FBH1 KO cells and FBH1 KO cells expressing wild-type FBH1 or E3 ligase-deficient FBH1 (*i*.*e*., FBH1-F266A/P267A [9]) and transfected with negative control (Ctrl) or RAD54L siRNA (n=2; 219-401 fiber tracts analyzed; ns=not significant, ****p < 0.0001; Kruskal-Wallis test followed by Dunn’s multiple comparisons test).

### The FBH1 E3 ligase mutant is proficient in RF reversal without RAD54L

U2OS FBH1 KO cells complemented with an E3 ligase-deficient FBH1 variant (FBH1-UbiM) are fully proficient in RF reversal [9, 23], suggesting that FBH1’s E3 ligase activity is not required for efficient RF reversal. To determine whether FBH1-UbiM cells rely on RAD54L for RF reversal, we depleted RAD54L in FBH1 KO, FBH1 KO+FBH1, and FBH1 KO+FBH1-UbiM cells (Fig. S1C) and measured IdU tract lengths, an indicator of RF restraint [33], using the DNA fiber assay under a low-dose HU treatment (Fig. 1C). Consistent with our previous observations in HeLa cells [20], IdU tracts were significantly shorter in FBH1 KO cells depleted of RAD54L than in control-depleted cells (Figs.1D, S1D). These results suggest that, under low concentrations of HU, the release of RF restraint in the absence of FBH1 relies on RAD54L. In addition, RF restraint was compromised in FBH1 wild type U2OS cells upon RAD54L knockdown, again in agreement with our earlier findings [20]. In contrast, FBH1 KO+FBH1-UbiM cells retain RF restraint in the absence of RAD54L with no significant difference relative to control-depleted FBH1 KO+FBH1-UbiM cells (Figs. 1D, S1D). Collectively, these results suggest that RF reversal can proceed efficiently in FBH1-UbiM cells without a requirement for RAD54L.

### RAD54L and FBH1 associate in human cells and complex formation is enhanced upon treatment of cells with HU

To further investigate the cooperation between wild type FBH1 and RAD54L in RF reversal, we transiently expressed EGFP-FBH1 in RAD54L KO cells and in RAD54L KO cells complemented with wild type RAD54L-HA [31]. Co-immunoprecipitation analyses revealed that both EGFP-FBH1 and RAD51 are present in anti-HA complexes isolated from whole-cell lysates (Fig. 2A, lane 3). Consistently, endogenous RAD54L was detected in anti-GFP precipitates from HeLa cells transiently expressing peGFP-FBH1 (Fig. 2B, lane 3), but not in anti-GFP precipitates from RAD54L KO cells (Fig. 2B, lane 4).

**Figure 2.**
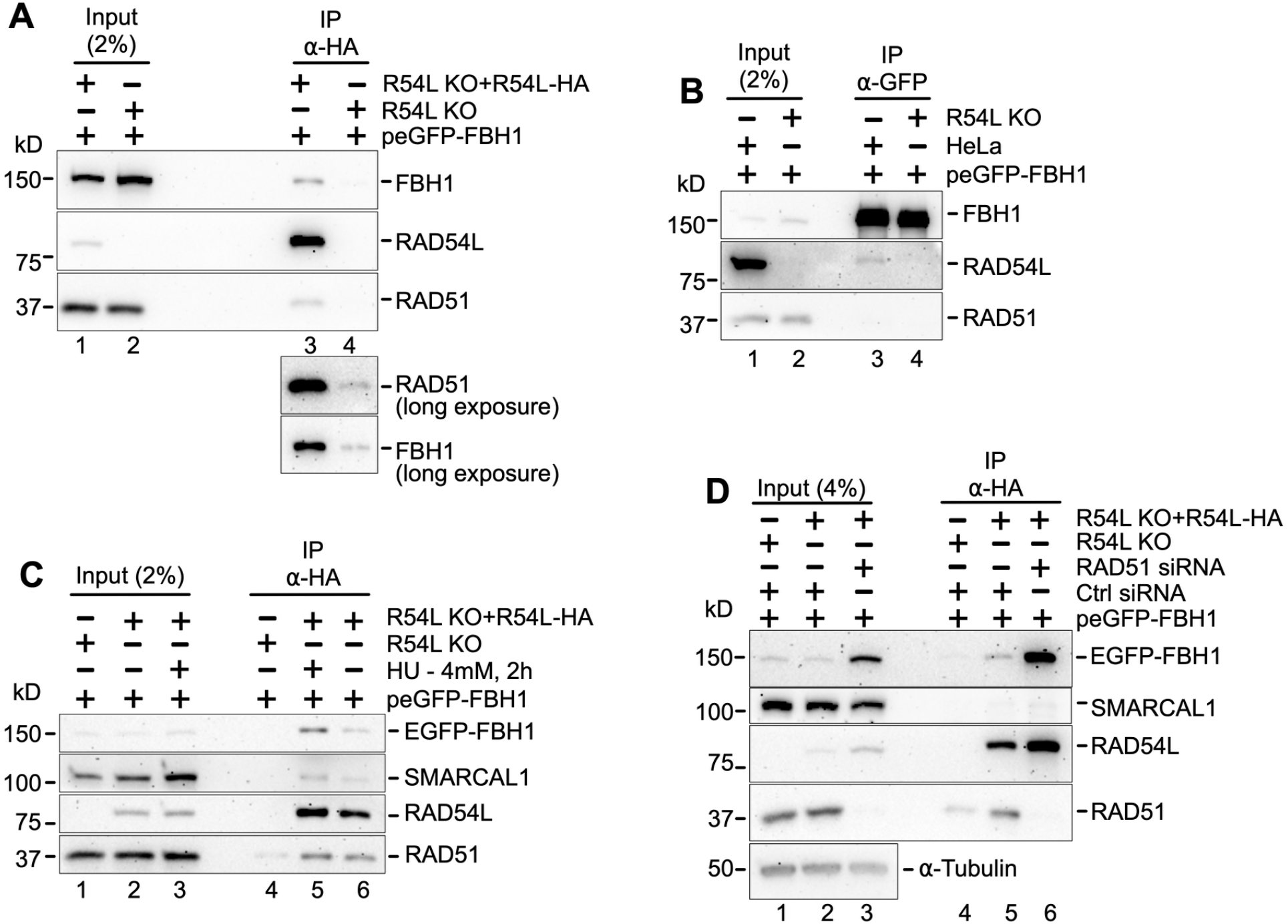
RAD54L and FBH1 co-precipitate in human cells. (**A**) Western blots to show that transiently expressed EGFP-FBH1 co-precipitates in anti-HA complexes in whole cell lysates prepared from RAD54L KO+RAD54-HA cells (lane 3). RAD51: positive control [19]. Lane 4: negative control. EGFP-FBH1 does not bind non-specifically to anti-HA resin on RAD54L KO cells. IP: immunoprecipitation. (**B**) Western blots to show that endogenous RAD54L co-precipitates in anti-EGFP complexes of whole cell lysates prepared from HeLa cells transiently transfected with peGFP-FBH1 (lane 3). (**C**) Western blots to show that transiently expressed EGFP-FBH1 is recruited to anti-HA protein complexes prepared from RAD54L KO+RAD54L-HA cells after a 2-h incubation in 4 mM HU (lane 5). Lane 6: cells not treated with HU. Lane 4: Neither EGFP-FBH1 nor RAD51 co-precipitate in anti-HA protein complexes prepared from whole cell lysates of RAD54L KO cells transiently transfected with peGFP-FBH1. (**D**) Western blots to show that more EGFP-FBH1 is present in anti-HA complexes prepared from whole cell lysates of RAD54L KO+RAD54L-HA cells transfected with RAD51 siRNA (lane 6) than in RAD54L KO+RAD54L-HA cells transfected with Ctrl siRNA (lane 5).

Following HU treatment of RAD54L KO+RAD54L-HA cells, EGFP-FBH1 was enriched in anti-HA precipitates (Fig. 2C, compare lanes 5 and 6), indicating that RAD54L-FBH1 complex formation is stimulated by RF stalling. Since both RAD54L and FBH1 are known to directly interact with RAD51 [19, 28], we next assessed RAD54L-FBH1 complex formation under conditions of RAD51 depletion. Notably, greater amounts of EGFP-FBH1 were detected in anti-HA precipitates from RAD54L KO+RAD54L-HA cells transfected with RAD51 siRNA compared to cells transfected with control siRNA (Fig. 2D, lanes 5 and 6). These results indicate that RAD54L-FBH1 complex formation is unlikely to be mediated by RAD51. Moreover, these results suggest that the RAD54L-FBH1 complex in cells is enhanced in the absence of RAD51.

### FBH1 predominantly associates with the RAD54L-Lobe1 domain

To further define how FBH1 interacts with RAD54L, we adopted an approach previously used to map the NUCKS1 interaction on RAD54L [30]. EGFP-FBH1 was transiently expressed with either HA-tagged full-length RAD54L or individual RAD54L domains (Fig. 3A) in RAD54L KO cells, followed by immunoprecipitation using anti-HA resin. These experiments revealed that EGFP-FBH1 associates most strongly with the RAD54L-Lobe 1 domain (Fig. 3B, lane 10).

**Figure 3.**
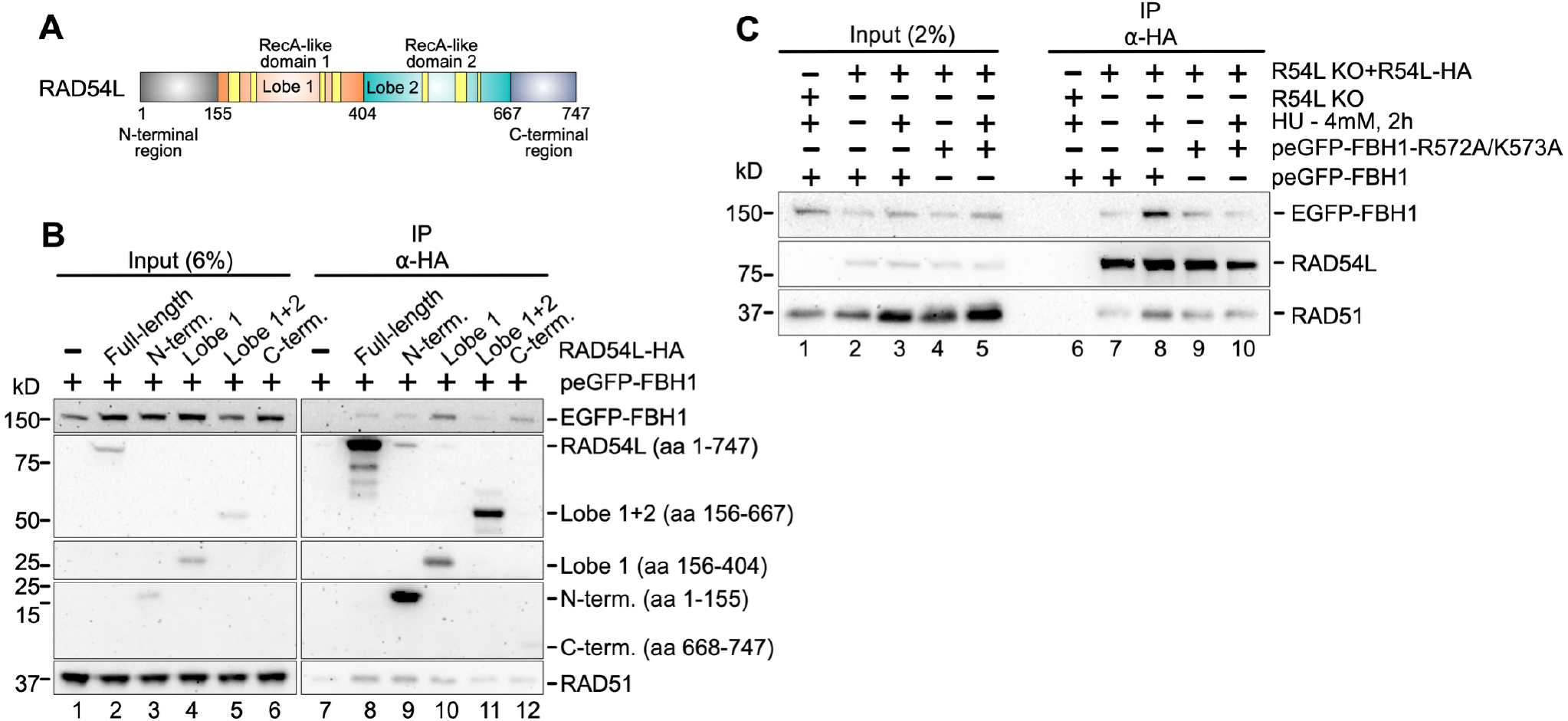
FBH1 predominantly associates with RAD54L-Lobe 1 in cells, and their association at stalled replication forks is dependent on the identified [24] FBH1 fork junction binding motif. **(A)** Schematic of the human RAD54L protein depicting the N-terminal domain, the two RecA-like domains (lobes 1 and 2), and the C-terminus. The seven conserved ATPase motifs are shown in yellow. (**B**) Western blots to show that transiently expressed EGFP-FBH1 co-precipitates predominantly with RAD54L-Lobe 1 (lane 10) in whole cell lysates of transiently transfected HeLa RAD54L KO cells. Lane 7: EGFP-FBH1 does not bind to the anti-HA resin non-specifically. (**C**) Western blots to show that transiently expressed EGFP-FBH1 is enriched in anti-HA complexes prepared from chromatin-bound protein fractions of RAD54L KO+RAD54L-HA cells after 2-h treatment with 4 mM HU (compare lanes 7 and 8). In contrast, an FBH1 mutant (R572A/K573A) with an abrogated RF junction binding motif [24] is not enriched after treatment of cells with HU (compare lanes 9 and 10).

We next used site-directed mutagenesis to disrupt the reported RF junction-binding motif in FBH1 (R447/K448 in isoform 4 [24], corresponding to R572/K573 in isoform 1). We first confirmed that EGFP-FBH1 and EGFP-FBH1-R572A/K573A were expressed at comparable levels in transiently transfected cells (Fig. S2A). FBH1 KO cells expressing FBH1-R572A/K573A exhibited defective RF restraint (Fig. S2B-2D), as reported [24], indicating that the EGFP-tagged FBH1 variants behave as expected. RAD54L KO+RAD54L-HA cells were then transfected with either peGFP-FBH1 or the R572A/K573A mutant, treated with HU (4 mM, 2 h) or left untreated, and chromatin extracts were subjected to anti-HA immunoprecipitation. EGFP-FBH1, but not the R572A/K573A mutant, was enriched in anti-HA precipitates from HU-treated cells (Fig. 3C, compare lane 8 to lane 10). These findings suggest that FBH1 associates with RAD54L at stalled RFs in cells, and that this interaction depends on an intact RF junction-binding motif in FBH1.

### Purified FBH1 and RAD54L physically interact

To determine whether the interaction between FBH1 and RAD54L is direct, we purified both proteins (Figs. S3A-S3C) and performed reciprocal pull-down assays using anti-FLAG M2 and anti-GST resins. The results show that FBH1 directly binds RAD54L-FLAG in anti-FLAG pull-downs (Fig. 4A, lane 4). Conversely, RAD54L specifically co-precipitated with GST-FBH1 (Fig. 4B, lane 4), further confirming a direct interaction between the two proteins.

**Figure 4.**
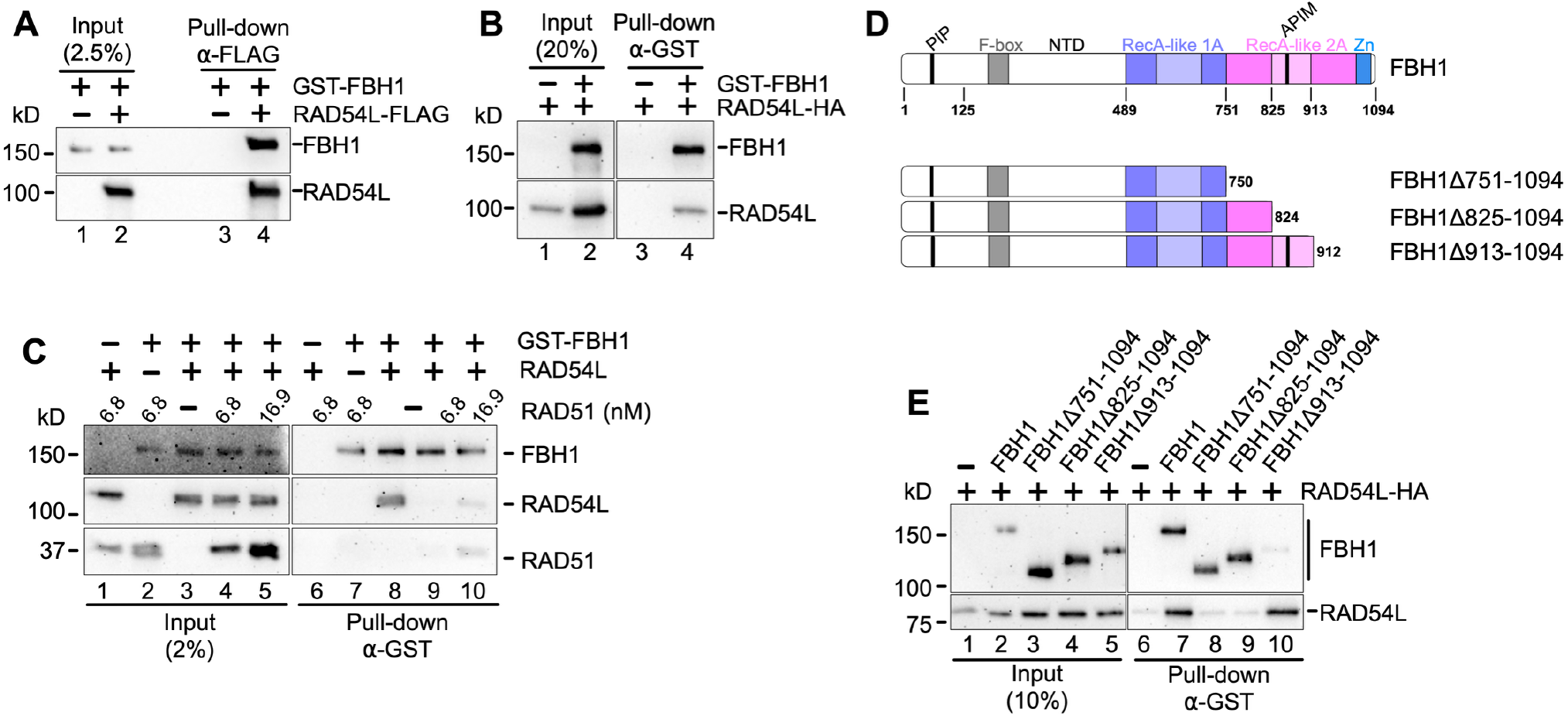
RAD54L and FBH1 directly interact. (**A**) Western blots to show that purified RAD54L-FLAG and GST-FBH1-His_6_ directly interact in a FLAG pull-down assay (lane 4). FBH1 does not bind to the FLAG resin (lane 3). (**B**) Western blots to show that purified GST-FBH1-His_6_ and RAD54L-HA directly interact in a GST pull-down assay (lane 4). RAD54L does not bind to the GST resin (lane 3). (**C**) Western blots to show that RAD54L forms a stable complex with FBH1 in GST pull-down assays (lane 8). Incubation of RAD51 (6.8 and 16.9 nM) with both FBH1 and RAD54L interferes with the FBH1-RAD54L complex (lanes 9 and 10). (**D**) Top: Schematic of the 1094 amino acid human FBH1 protein (isoform 1) depicting the N-terminal domain and two RecA-like domains. Bottom: Schematics of the three FBH1 truncations tested. (**E**) Western blots to show that RAD54L interacts with the 2B subdomain (residues 751-912; rose) in the FBH1 RecA-like 2A domain (pink).

We next performed competitive pull-down assays to determine whether purified RAD51 (Fig. S3D) interferes with the FBH1-RAD54L interaction. The results show that addition of excess RAD51 (6.8 and 16.9 nM) disrupted the FBH1-RAD54L complex (Fig. 3C, compare lane 8 to lanes 9 and 10). The results also show that, under these conditions, FBH1 and RAD51 do not co-precipitate on GST resin (Fig. 4C, lane 7). Together, these results suggest that RAD54L preferentially associates with RAD51 over FBH1.

To map the interaction interface of RAD54L on FBH1, we generated structural models of the FBH1-RAD54L complex using AlphaFold 3 (AF3) [34, 35]. AF3 output models were ranked according to per-residue and interface confidence scores. The top-ranked model (Model 0) was selected for subsequent analyses in UCSF ChimeraX and superimposed onto the published cryo-EM structure of the SCFFBH1 complex at a three-way model RF substrate (PDB ID: 9XZJ; [24]) (Fig. S3E). Residues located at the predicted RAD54L-FBH1 interface within 3.5 Å were identified from the structural model (Table S1; Fig. S3F), revealing that RAD54L contact sites likely reside within residues 785–793 in FBH1 isoform 4, corresponding to residues 910–918 in isoform 1 (for schematic see Fig. 4D).

To validate these predictions experimentally, we generated and purified three truncated FBH1 variants: FBH1Δ751–1049 and FBH1Δ825–1049, both lacking the predicted RAD54L interacting region, and FBH1Δ913–1049, which retains most of the predicted contact residues (Fig. 4D). Full-length FBH1 and the FBH1 truncations were tested for RAD54L binding using GST pull-down assays. RAD54L co-purified with full-length FBH1 and with FBH1Δ913–1049 (Fig. 4E, lanes 7 and 10, respectively). In contrast, RAD54L failed to co-precipitate with either FBH1Δ751–1049 or FBH1Δ825–1049 (Fig. 4E, lanes 8 and 9). In agreement with our AF3 structural predictions, these findings suggest that a key RAD54L interaction motif resides within FBH1 residues 825–912. Together with the AF3-predicted contact sites (Table S1), these results identify residues I910, E911, and D912 as part of a critical FBH1 motif required for complex formation with RAD54L.

### Purified FBH1 and RAD54L functionally cooperate in reversal of a model RF

Both FBH1 and RAD54L are individually capable of catalyzing the reversal of model RFs in biochemical reconstitution assays [23, 36]. However, their coordinated activity in this process, previously observed in cells [20], had not been examined *in vitro*. To address this, we generated a preferred RF substrate for SCFFBH1 [24], a model RF containing an 8-bp gap on the lagging strand immediately adjacent to the junction (Fig. 5A). In addition, a 2-bp gap was introduced on the leading strand adjacent to the junction to enhance FBH1 binding [24]. Reversing the RF with a −8 bp lagging strand gap and a −2 bp leading strand gap generates both the parental DNA duplex and a nascent DNA duplex containing a 6-bp 3′ ssDNA overhang, which can be further unwound by the FBH1 helicase (Figs. 5A, S4A, S4B).

**Figure 5.**
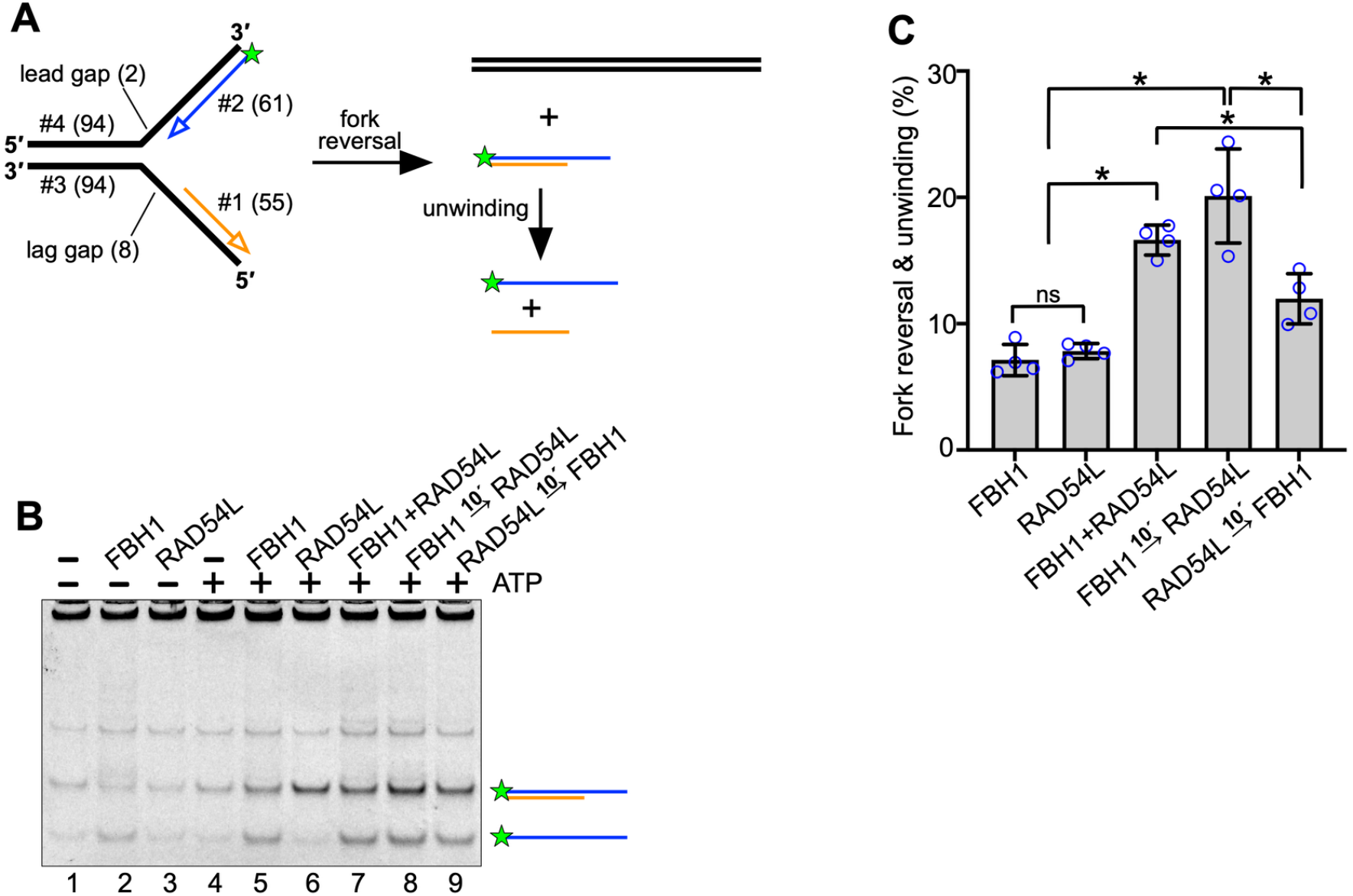
RAD54L and FBH1 cooperate in reversal of a model *RF in vitro*. (**A**) Schematic of the model RF used in (B) and (C) and of the reversal and unwinding products. The Cy5 labeled oligonucleotide (#2; leading strand) is depicted by a green asterisk. (**B**) Representative gel image of the *in vitro* fork reversal assays with FBH1 and/or RAD54L. Gel auto-adjusted for visualization. (**C**) Quantification of the percentage of reversal and unwinding products. n=4; ns, not significant; *, p < 0.05; two-tailed t-test.

We performed the reactions in the presence or absence of ATP using purified FBH1 (59 nM) and/or RAD54L (135 nM). Individually, RAD54L and FBH1 showed similar activity in promoting RF reversal and reversal plus associated unwinding, respectively (Fig. 5B, lanes 5 and 6; Fig. 5C). In contrast, simultaneous addition of both proteins markedly increased product formation (Fig. 5B, lane 7; Fig. 5C). Furthermore, order-of-addition experiments revealed that reversal efficiency was greater when FBH1 was added prior to RAD54L (Fig. 5B, lane 8; Fig. 5C). These results strongly suggest that FBH1 initiates the reaction prior to engaging RAD54L and provide the first mechanistic insight into the sequential coordination of FBH1 and RAD54L activities during RF reversal.

## DISCUSSION

Replication fork (RF) reversal serves as a protective mechanism that stabilizes stalled RFs to allow repair of DNA lesions without causing chromosomal breakage. RF reversal is a highly coordinated process and is mediated by several RF remodeling enzymes including SMARCAL1, ZRANB3, HLTF, and the 3′-5′ FBH1 helicase [4]. SMARCAL1, ZRANB3, and HLTF act coordinately in RF reversal, as inferred from nascent strand degradation assays [9]. In contrast, FBH1 operates in a distinct reversal pathway [9] and utilizes a fundamentally different molecular mechanism, as recently demonstrated [24].

In our previous study, we showed that RAD54L BM activity is selectively required in the FBH1-dependent reversal pathway [20]. Consistent with these findings, we show here that a separation-of-function RAD54L mutant (RAD54L-BM), deficient in BM but proficient in the stimulation of RAD51-mediated strand exchange [32], prevents nascent strand degradation in the absence of 53BP1, a RF protection factor specific to the FBH1 pathway [9]. Importantly, this RAD54L-BM mutant does not prevent nascent strand degradation in the SMARCAL1/HLTF/ZRANB3 pathway, as we have previously shown [20]. Together, these results corroborate our earlier findings that identified that RAD54L BM activity is selectively required in the FBH1 pathway [20].

Here, we provide new biochemical evidence supporting the coordinated action of RAD54L and FBH1 in RF reversal. We demonstrate that RAD54L and FBH1 physically and functionally interact. In cells, the two proteins associate at stalled RFs, and complex formation is dependent on the identified RF junction-binding motif in FBH1 [24]. These results suggest that FBH1 localizes to RF junctions prior to RAD54L, facilitating complex assembly. In accord, biochemical reconstitution assays show that RF reversal is more efficient when FBH1 is added to the reaction first. These results suggest that the FBH1 helicase initiates the reversal reaction and unwinds the lagging strand, as proposed [24, 29], thereby creating a substrate for RAD54L BM. Such an intermediate may resemble a partial Holliday (PX) junction, a preferred substrate for RAD54L BM activity [37].

Surprisingly, in contrast to FBH1 wild-type cells, RAD54L does not appear to contribute to RF reversal in FBH1 E3 ligase mutant cells, as shown here. These findings could suggest that, in the presence of this FBH1 mutant, elevated RAD51 protein [25] may associate with stalled RFs, enabling the formation of the 4-way junction and catalysis of BM without RAD54L. Notably, purified RAD51 is capable of catalyzing BM *in vitro* through a very distinct mechanism that relies on repetitive cycles of RAD51 polymerization and dissociation [38]. As such, sufficient nuclear RAD51 may be critical for the repetitive nature of this process in cells. In the absence of the FBH1 E3 ligase, RAD51 no longer is targeted for ubiquitylation [25], potentially enabling RAD51’s continuous reassociation with the reversing RF. In contrast, under wild type conditions, E3 ligase-proficient FBH1 ubiquitylates RAD51 and, thereby, limits RAD51 association with the chromatin [25, 27, 28], which then may necessitate RAD54L BM activity in the FBH1-mediated RF reversal pathway.

However, our results in E3 ligase-deficient FBH1 cells raise a second possibility that may explain why RAD54L is not needed for the catalysis of RF reversal by this FBH1 mutant. FBH1 is an F-box adaptor protein in the SCF (SKP1-CUL1-ROC1-F-box protein; SCFFBH1) complex [22]. FBH1 is linked to the core complex through the interaction between SKP1 and the FBH1 F-box domain [39]. Ubiquitin is transferred from the ubiquitin-conjugating enzyme (E2) onto the substrates recruited by FBH1. However, E3 ligase-deficient FBH1 contains mutations in key residues within the F-box (F266A/P267A) and does not bind to SKP1 in cells [22, 27]. Moreover, SKP1 bridges the interaction between FBH1 and CUL1 and ROC1 [22, 40]. Hence, the E3 ligase deficient FBH1 mutant additionally is unlikely to associate with CUL1 or ROC1 in cells. Taken together, the results from these earlier studies suggest that the SCFFBH1 complex likely is not assembled in FBH1 KO cells that express E3 ligase-deficient FBH1. As these mutant cells retain wild type proficiency in RF reversal, FBH1, in contrast to SCFFBH1, may be able to catalyze the reversal reaction without RAD54L. As both helicase and ATPase activities in the SCFFBH1 complex are indistinguishable from those in FBH1 alone [26], we speculate that FBH1 alone and without RAD54L may function at a subset of stalled RFs. These could be RFs with the CMG complex retained [21], which is predicted to preclude binding of SCFFBH1 complex to the RF junction [24].

A recent cryo-EM structure of SCFFBH1 bound to a three-way DNA substrate reveals that FBH1 promotes annealing of parental DNA strands through a mechanism distinct from that of SMARCAL1, HLTF, and ZRANB3 [24]. Whereas these enzymes drive RF reversal by pushing from the parental duplex toward the RF, SCFFBH1operates from behind the RF, pulling the lagging strand template inward [24]. However, this model does not fully explain how the characteristic four-way junction of a reversed RF is formed or how base pairing between nascent strands is established. Unlike SMARCAL1, HLTF, and ZRANB3 [24, 41, 42], SCFFBH1 lacks intrinsic strand annealing activity [24], highlighting a gap in our understanding of the additional factors required for the FBH1-mediated pathway.

Like SMARCAL1, HLTF, and ZRANB3, RAD54L translocates along dsDNA in an ATP-dependent manner [4, 43, 44]. It also shows a strong preference for binding PX junctions, and to a lesser extent for other branched DNA structures such as three-way and Holliday junctions [37]. During RF reversal, RAD54L may be positioned at the RF junction through the interaction with FBH1. Using FBH1 variants with different deletions in the RecA-like 2A domain, we identified the FBH1 2B subdomain (residues 825-912) as critical in mediating the interaction with RAD54L. Moreover, AF3 structural predictions narrowed the contacting motif to three C-terminal residues (I910, E911, and D912) in the 2B region. Of note, the SCFFBH1 structure shows that the FBH1 2B subdomain does not contact DNA [24], suggesting that-in this presumably active state-the FBH1 2B subdomain has the ability to interact with other partners. As some SF1 helicases are activated through protein-protein interactions within their 2B subdomains [45, 46], we speculate that the 2B subdomain may be a critical regulatory domain in FBH1 as well, and that RAD54L may activate FBH1 in RF reversal through its interaction with the FBH1-2B region. This would be similar to the activation of the UvrD helicase by MutL in DNA mismatch repair, which binds to the UvrD 2B subdomain [46]. This proposal is also substantiated by our results obtained in biochemical reconstitution assays, in which FBH1 is significantly more active in RF reversal when RAD54L is present. In contrast, adding RAD54L to the reaction first reduces reversal product formation, suggesting that FBH1 is less apt to stimulate RF reversal by RAD54L.

In our preferred model of FBH1 and RAD54L cooperativity at RFs, FBH1 initiates the reversal process and then dissociates from the RF, enabling a conformational change in RAD54L for further engagement with the RF (Fig. 6). FBH1’s prolonged residence at the RF junction may sterically hinder the formation of the four-way junction structure and BM [24]. Like SMARCAL1, HLTF, and ZRANB3, RAD54L then may promote RF remodeling by pushing from the parental duplex toward the RF.

**Figure 6.**
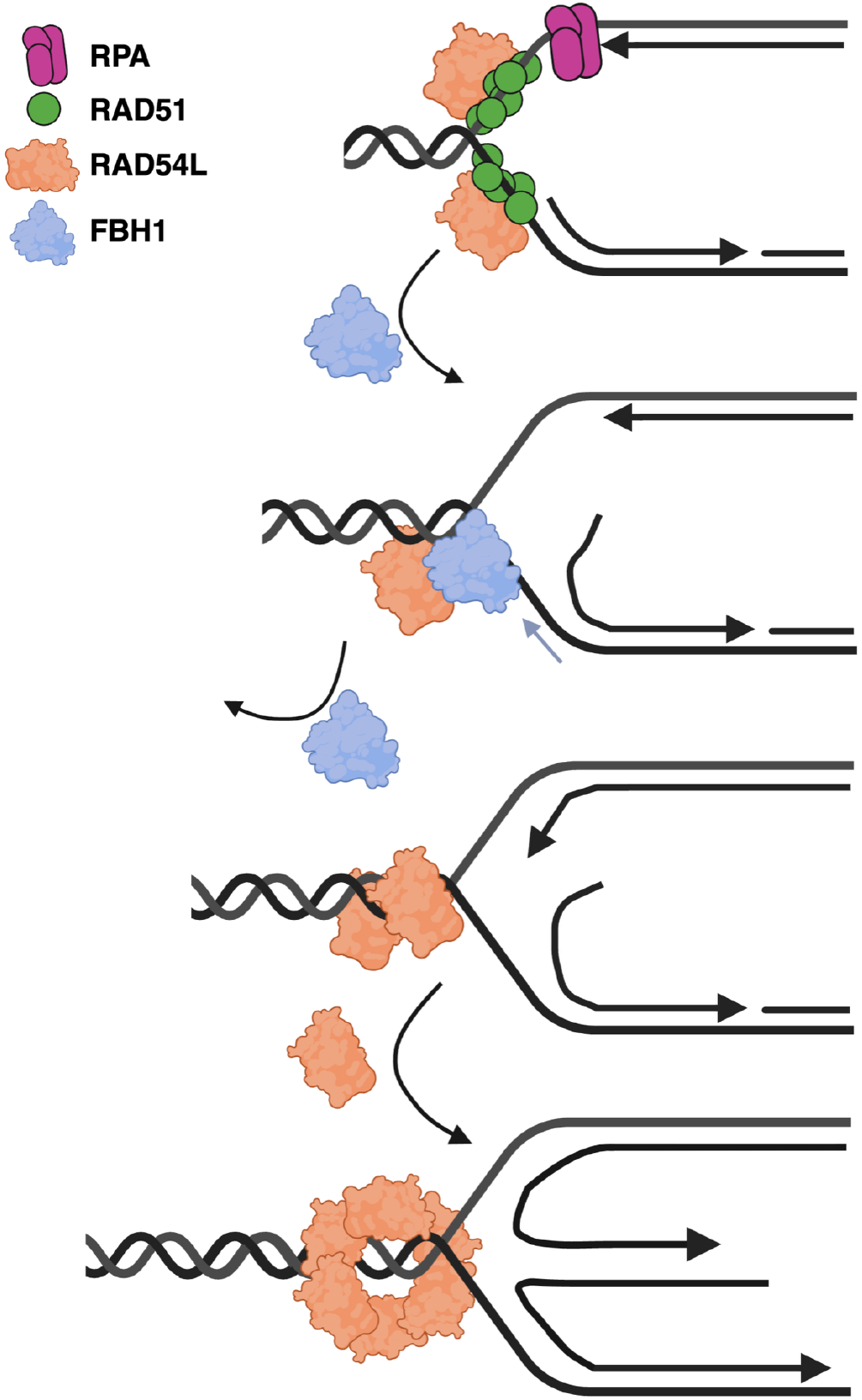
Model to illustrate how FBH1 and RAD54L orchestrate replication fork reversal. FBH1 binds to the RF junction first (a.) and initiates RF reversal through moving the lagging strand template towards the RF junction, extending the annealed portion of the parental DNA strands, as predicted [24]. FBH1 also initiates unwinding of the lagging strand [24, 29]. FBH1 recruits RAD54L to the RF junction (b.) and dissociates from the RF (c.) to allow assembly of oligomeric RAD54L for branch migration (d.). How the nascent strands are annealed to facilitate the formation of the 4-way junction remains unclear.

Downstream of SMARCAL1, ZRANB3, and HLTF, TOP2A initiates more extensive RF reversal, via recruitment of the PICH translocase, which, like RAD54L [36, 37, 47, 48], can BM four-way DNA junctions [49]. These observations led to the proposal that RF reversal in the SMARCAL1/ZRANB3/HLTF pathway involves the sequential action of at least two enzymatic activities [49]. First, RF reversal activity leads to the conversion of stalled RF fork into a four-way junction. Second, BM activity converts the four-way junction into more extensively reversed RFs [49]. How exactly the four-way junction is created in the FBH1/RAD54L pathway remains unclear. Moreover, if more extensive RF reversal in the FBH1/RAD54L pathway requires another enzyme capable of BM, such as PICH helicase, has not been tested yet. As RAD54L is one of the most efficient BM enzymes in eukaryotic cells and capable of bypassing long stretches of sequence heterology [50], RAD54L may be the only enzyme needed in the catalysis of BM in the FBH1 RF reversal pathway.

## MATERIALS AND METHODS

### Cell lines, transfections, siRNAs, and expression of ectopic proteins in human cells

HeLa cells were obtained from ATCC and maintained as recommended. HeLa cells that are knockout (KO) for *RAD54L*, and HeLa *RAD54L* KO cells ectopically expressing RAD54L were maintained as described previously [20, 30, 31]. U2OS *FBH1* KO cells [9] were a gift from Dr. David Cortez (Vanderbilt) and grown in Dulbecco’s modified Eagle’s medium (DMEM) with 10% fetal bovine serum.

The negative control siRNA (Ctrl) and siRNAs targeting RAD54L and RAD51 were described earlier [20, 30] and were obtained from IDT (Table S2). SiRNA forward transfections with Lipofectamine RNAiMAX (Thermo Fisher Scientific) were performed on two consecutive days with a concentration of 20 nM for each siRNA. Transfection of cells with plasmids to express EGFP-FBH1 (peGFP-FBH1) or RAD54L-HA and RAD54L truncations [30] were performed with Lipofectamine 2000 (Thermo Fisher Scientific) and as described [30].

The human RAD54L deletion constructs were described previously [30]. To generate peGFP-FBH1, full-length FBH1 was amplified from pLNCX-3XHA-FBH1 [9] using the primer pairs listed in Table S3 and cloned from *Xho*I to *Kpn*I into pEGFP-C1 (Clontech). To generate peGFP-FBH1-R572A/K573A, mutations were generated in peGFP-FBH1 using the primers listed in Table S4 and Q5 Site-Directed Mutagenesis Kit (New England Biolabs).

The vectors containing the C-terminally HA-tagged full-length human FBH1 or F-box mutant FBH1-F266A/P267A were gifts from Dr. David Cortez [9]. FBH1-HA cDNAs were amplified from pLNCX-3XHA-FBH1 (or - F266A/P267A) using the primers listed in Table S3, cloned from *KpnI* and *XhoI* into pENTR1A (Invitrogen), and transferred into pLentiCMV/TO DEST#2 [51] with Gateway LR Clonase II (Invitrogen). Lentivirus was generated in HEK293FT cells (Invitrogen) and used to transduce U2OS FBH1 KO cells in 6 μg/ml polybrene, as described [51].

### Western blot analysis

Western blotting was performed according to our established protocol [52]. The following primary antibodies were used: α-RAD54L (F-11; sc-374598; Santa Cruz Biotechnology; 1:1000), α-RAD51(Ab-1; EMD Millipore; 1:4000), α-FBH1 (sc-81563;1:2000), α-SMARCAL1 (sc-166209; 1:300), α-HA (16B12; 901533; BioLegend), α-α-Tubulin (DM1A; Santa Cruz Biotechnology; 1:1000), α-GFP (50430-2-AP; Proteintech; 1:8,000), α-53BP1 (A300-272A; Bethyl Laboratories; 1:30,000). HRP-conjugated goat anti-rabbit or goat anti-mouse IgG (Jackson ImmunoResearch; 1:10,000) secondary antibodies were used. Western blot signals were acquired with a Chemidoc XRS+ gel imaging system and ImageLab software version 6.1 (BioRad).

### DNA fiber assay

The DNA fiber assay was used essentially as described earlier [20, 53]. Briefly, cells were pulse labeled with 25 µM CldU for 20 min followed by 250 µM IdU containing 25 µM hydroxyurea (Sigma) for 30 min. Only IdU tracts directly following a CldU tract were measured. To assess nascent strand degradation in the absence of RF protection, cells were pulse labeled with 25 µM CldU for 20 min and then 250 µM IdU for 20 min before a 5-h treatment with 4 mM HU.

### Expression vectors for protein purification from Sf9 cells

To generate human RAD54L-HA for expression in Sf9 insect cells, RAD54L-HA in pENTR1A (Invitrogen), described previously [31], was used in the BaculoDirect™ system (Invitrogen). RAD54L-HA was transferred from pENTR1A into BaculoDirect™ Linear DNA through a Gateway® Clonase II LR reaction and transfected into Sf9 insect cells (Invitrogen) for virus production following a standard protocol (Invitrogen).

To generate full-length GST-FBH1-His_6_, FBH1 was amplified from pLNCX-3XHA-FBH1 WT [9] using the primer pairs listed in Table S3 and cloned from *SalI* to *NotI* into pGEX-6P-1 (Cytiva). GST-FBH1-His_6_ DNA was then amplified from pGEX-6P-1-FBH1 using the primer pairs listed in Table S3 and cloned from *KpnI* to *NotI* into pENTR1A (Invitrogen). To generate GST-FBH1 truncations (FBH1Δ751–1049, FBH1Δ825–1049, and FBH1Δ913–1049), STOP codons were introduced in pENTR1A-GST-FBH1-His_6_ using the primers listed in Table S4 the Q5 Site-Directed Mutagenesis (New England Biolabs). All ORFs were transferred from pENTR1A into BaculoDirect™ Linear DNA *via* Gateway® Clonase II LR recombination and transfected into Sf9 insect cells (Invitrogen) for virus production following a standard protocol (Invitrogen)

### Recombinant protein purification

#### His6-Smt3-RAD51 was expressed and purified as described [54-56]

Untagged RAD54L was purified as previously reported [56]. Briefly, 1.5 ⨉ 109 Sf21 cells were infected with recombinant baculovirus (high-titer P3 viral stock; multiplicity of infection = 5 PFU/cell) and incubated for 48 h at 27 °C. Harvested cells (13.6 g) were resuspended in 100 mL lysis buffer. Clarified lysate was applied to a 30 mL Q-Sepharose column. The flow-through was collected, loaded onto a 10 mL glutathione-Sepharose column, and eluted with buffer containing 20 mM glutathione. The protein was concentrated to 1 mL in a dialysis bag surrounded by precooled solid polyethylene glycol (Mr 20,000 Da). Thrombin was then added (50 μg per mg of RAD54L), and the sample was incubated for 5 h at 4 °C. The cleaved protein was loaded onto a 60 mL Superdex-200 column. Fractions free of ssDNA-specific nuclease activity were pooled, diluted to 100 mM KCl, loaded onto a 1 mL Resource-S column and eluted with a 100-500 mM KCl gradient. Fractions containing untagged RAD54L were pooled and dialyzed overnight against 20 mM Tris-HCl, pH 7.5, 400 mM KCl, 30% glycerol, 1 mM DTT, then frozen and stored at -80 °C.

#### FLAG-tagged RAD54L was purified as previously reported [30, 57]

For expression of RAD54L-HA in Sf9 cells, 1.5 ⨉ 108 Sf9 cells were infected with a high-titer P3 viral stock using a multiplicity of infection of 10 PFU/cell and incubated at 28 °C for 48 h. Harvested cells were resuspended in 10 ml lysis buffer (50 mM Tris pH 7.5, 200 mM KCl, 2 mM EDTA, 10% glycerol, 10 mM β-mercaptoethanol (BME), and 0.5% NP-40) containing EDTA-Free Protease inhibitor (Pierce) and incubated on ice for 15 min before 4 cycles of sonication (20 s on and 40 s off). After centrifugation (24,000 ⨉ g for 15 min at 4 °C), the clarified lysate was diluted 1:5 with wash buffer (20 mM Tris pH 7.5, 150 mM KCl, 0.1 mM EDTA, and 0.05% NP-40) and incubated with anti-HA affinity matrix (Roche) on a rotator at 4 °C for 2 h with end-over-end mixing. Unbound protein was collected after centrifugation at 300 ⨉ g for 1 min, and the resin was washed three times with wash buffer. Bound proteins were eluted with 2 mg/ml HA peptide (Thermo Fisher Scientific) in 3 consecutive 15 min elutions on ice.

For expression of GST-FBH1-His_6_ and FBH1 variants, 1.5 ⨉ 108 Sf9 cells were infected with a high-titer P3 viral stock using a multiplicity of infection of 10 PFU/cell and incubated at 28 for 48 h. Harvested cells were resuspended in 23 ml GST Lysis Buffer (50 mM Tris-HCl pH 8.0, 300 mM NaCl, 1 mM EDTA, 10% glycerol, 2 mM DTT, 0.1% TritonX-100) with EDTA-Free Protease inhibitor (Pierce) and incubated on ice for 15 min before 4 cycles of sonication (20 s on and 40 s off). After centrifugation (24,000 g for 15 min at 4 °C), the clarified lysate was incubated with 750 µL packed glutathione agarose (Sigma) for 16 h. Unbound protein was collected following centrifugation at 300 ⨉g for 3 min, and the resin was washed four times with 10 ml GST lysis buffer. Washed resin was incubated with 100 µl GST elution buffer (40 mM Tris-HCl pH 8.0, 50 mM NaCl, 10% glycerol, 10 mM glutathione) for 15 minutes on ice. The elution step was repeated twice more with a total of three 100 µl elutions collected.

### GST Pulldowns

Two hundred ng GST-FBH1-His_6_ (3.3 nM) or FBH1 variants (4.6-3.9 nM) alone or with 60 ng RAD54L-HA (1.8 nM) was incubated in 400 µl binding buffer (10 mM Tris-HCl pH 8.0, 150 mM NaCl, 1mM EDTA, 0.1% TritonX-100, 1 mM DTT, 1 mM MgCl_2_) at 4 for 4 h with gentle agitation in the presence of 250 U benzonase (Sigma). Next, glutathione agarose (Sigma) was equilibrated in binding buffer, and 20 µl packed resin was added to each sample and incubated overnight at 4 °C with end-over-end mixing. Unbound protein was removed from the resin by centrifugation at 300 ⨉g for 5 min followed by four washes in 500 µl of binding buffer. Bound protein was eluted during two consecutive 15-min incubations in 25 µl GST elution buffer (40 mM Tris-HCl pH 8.0, 50 mM NaCl, 10% glycerol, 10 mM glutathione). Remaining protein bound to the resin was eluted in 25 µl of 2⨉ SDS-loading buffer and boiled at 95 °C for 5 min. Eluted protein was fractionated by SDS-PAGE, transferred to a PVDF membrane, and detected by Western blot.

For the competitive GST pulldowns shown in Figure 4C, 200 ng GST-FBH1-His_6_ (3.3 nM) was incubated with either 100 ng RAD51 (6.8 nM) or 60 ng RAD54L-HA (1.8 nM). Samples with RAD51 competitor included 200 ng GST-FBH1-His_6_ (3.3 nM), 60 ng RAD54L-HA (1.8 nM), and either 100 ng or 250 ng RAD51 (6.8 and 16.9 nM, respectively). All samples were incubated and processed as above.

### FLAG pulldown

FLAG pull-downs were performed as described with some modifications [30, 31]. Briefly, DYKDDDDK Fab-Trap® Agarose (ChromoTek) was equilibrated in binding buffer (50 mM Tris-HCl pH 7.5, 150 mM NaCl, 0.1% Triton X-100, and 100 μg/ml BSA). RAD54L-FLAG (80 nM) or no protein were incubated with the equilibrated resin at 4 °C for 1 h. Unbound protein was removed by centrifugation at 3,000 rpm for 3 min. GST-FBH1-His_6_ (80 nM) was added to the washed resin in 100 µl binding buffer and incubated at 4 °C for 1 h with gentle mixing in the presence of DNase I (1 U/µg protein). Unbound protein was removed, and the resin was washed three times with 200 µl binding buffer. Bound proteins were eluted in binding buffer containing 150 ng/μl 3X FLAG peptide (Sigma). Eluted protein was fractionated by 10% SDS-PAGE, transferred onto a PVDF membrane and detected by Western blot analysis.

### Anti-HA immunoprecipitations

HeLa *RAD54L* KO cells ectopically expressing RAD54L with a C-terminal HA tag [31] were transfected with peGFP-FBH1 plasmid (3.4 µg) and Lipofectamine 2000 (Thermo Fisher Scientific). For anti-HA immunoprecipitations of full-length RAD54L or RAD54L truncations, HeLa *RAD54L* KO cells [30, 31] were transfected with peGFP-FBH1 (240 ng) and/or plasmids containing either full-length RAD54L-HA or C-terminally HA-tagged RAD54L truncations (2.16 µg) [30]. Twenty hours after transfection, 3 ⨉ 106 cells were harvested and lysed in 35 µl lysis buffer (50 mM Tris-HCl, pH 7.5, 300 mM NaCl, and 0.5% NP-40 with protease and phosphatase inhibitors (Thermo Fisher Scientific). The lysates were centrifuged at 4 °C and adjusted to 150 mM NaCl and 0.05% NP-40, and 400 µg of total protein was incubated with 15 µl packed anti-HA affinity matrix (Roche) and 250 U benzonase (Sigma) on a rotator at 4 °C for 2 h with end-over-end mixing. Anti-HA resin was precipitated by centrifugation and washed 4⨉ in 500 µl of binding buffer. Bound proteins were eluted with 2⨉ SDS-loading buffer and boiled at 95 °C for 5 min. Eluted proteins (24 µl each) were detected by 10% SDS-PAGE and Western blot analysis.

For anti-HA immunoprecipitations following RAD51 depletion, HeLa *RAD54L* KO cells ectopically expressing RAD54L with a C-terminal HA tag [31] were transfected with 20 nM RAD51 or negative control siRNA and Lipofectamine RNAiMAX (Thermo Fisher Scientific) overnight before removing the transfection complex and replacing with fresh medium. Six hours later, the cells were transfected with 20 nM RAD51 or negative control siRNA, peGFP-FBH1 (570 ng), and Lipofectamine 2000 (Thermo Fisher Scientific) for four hours. On the next morning, 6 ⨉ 106 cells were harvested, lysed in 70 µl lysis buffer, and anti-HA immunoprecipitations were performed as described above. Here, 750 µg total protein were incubated with the anti-HA affinity matrix (Roche) to isolate proteins associated with RAD54L.

For anti-HA immunoprecipitation of chromatin bound proteins, HeLa *RAD54L* KO cells ectopically expressing RAD54L with a C-terminal HA tag [31] were transfected with peGFP-FBH1 or peGFP-FBH1-R572A/K573A (3.4 µg) and Lipofectamine 2000 (Thermo Fisher Scientific). On the next morning, the cells were treated with 4 mM hydroxyurea (Sigma) for 2 h. Around 2.5 ⨉ 107 cells were then trypsinized and harvested for subcellular protein fractionation using the Subcellular Protein Fractionation Kit for Cultured Cells (Thermo Fisher Scientific) as described by the manufacturer. Cytosolic, membrane-bound, soluble nuclear, and chromatin-bound protein fractions were isolated with the appropriate buffers and HALT protease and phosphatase inhibitors (Thermo Fisher Scientific). A total of 1.5 mg isolated chromatin-bound protein was diluted at a 1:9 ratio with binding buffer (25 mM Tris-HCl pH 7.5, 150 mM NaCl, and 0.05% NP-40) and incubated with 15 µl packed anti-HA affinity matrix (Roche) and 250 U benzonase (Sigma) on a rotator at 4 °C for 2 h with end-over-end mixing. Anti-HA resin was precipitated by centrifugation and washed 4⨉ in 500 µl binding buffer. Bound proteins were eluted with 2⨉ SDS-loading buffer and boiled at 95 °C for 5 min. Eluted proteins (24 µl each) were detected by 10% SDS-PAGE and Western blot analysis.

### Anti-GFP immunoprecipitation

HeLa *RAD54L* KO cells ectopically expressing RAD54L or *RAD54L* KO cells [30, 31] were transfected with peGFP-FBH1 (3.4 µg) and Lipofectamine2000 (Invitrogen). Twenty hours after transfection, 6 ⨉ 106 cells were harvested and lysed in 100 µl chilled lysis buffer (50 mM Tris-HCl, pH 7.5, 300 mM NaCl, and 0.5% NP-40) supplemented with EDTA-free protease inhibitor cocktail (Roche) and HALT phosphatase inhibitors (Thermo Fisher Scientific). Protein lysates were cleared by centrifugation at 4 °C and adjusted to 25 mM Tris-HCL, pH 7.5, 150 mM NaCl, and 0.05% NP-40. Samples were assembled with 1.75 mg total protein and 250 U benzonase (Sigma) and were incubated with 17 µl packed, equilibrated GFP-Trap Magnetic Particles (Chromotek) at 4 °C for 2 h with end-over-end mixing. Unbound protein was separated from GFP-Trap particles with a magnetic separation rack. GFP-Trap particles were washed four times with 500 µl binding buffer, and bound protein was eluted in 25 µl 2⨉ SDS-loading buffer and boiled at 95°C for 5 min. Eluted protein was fractionated by 10% SDS-PAGE, transferred to a PVDF membrane, and detected by Western blot.

### In vitro fork reversal assay

The oligonucleotides used to generate the fork reversal substrate are adapted from [36] and listed in Table S5. Forks were prepared by annealing the two arms separately. A 1.2-fold molar excess was used to anneal the unlabeled oligo #71 (3.6 µM) with the Cy5-labeled oligo #2 (3 µM) in 30 µl annealing buffer (100 mM NaCl, 50 mM Tris-HCl, pH7.5, 10 mM MgCl_2_, and 1 mM dithiothreitol (DTT)). Oligos #1 and #117 were annealed at equimolar ratio (3 µM) in a 30 µl annealing buffer. Reactions were heated to 94 °C for 3 min and cooled at 1 °C/min to 4 °C. To anneal the final fork, the unlabeled and labeled duplexes were annealed in a 2:1 molar ratio at 37 °C for 30 min. The fork reversal substrate was purified using 7% PAGE in 0.5⨉ TBE. The band was excised from the gel, the fork substrate was eluted into 0.5⨉ TBE, washed by sodium acetate precipitation and stored in TE, pH7.5, at -80 °C.

Fork reversal reactions were carried out in 17 µl total volume in fork reversal buffer (10 mM Tris-HCl, pH 7.5, 50 mM NaCl, 5 mM MgCl2, 100 µg/ml BSA, 2 mM ATP, and 1 mM DTT) [24]containing 6 nM fork substrate and proteins (as indicated) at 37 °C for 20 min. Reactions were terminated by the addition of 1 µl 10% SDS and 1 µl Proteinase K (≥ 500 U/ml; Sigma) at room temperature for 10 min. Three µl loading buffer (70% glycerol, 0.1% bromphenol blue) were added to the reactions and 3-5 µl of the sample were separated on a 9% 19:1 (acrylamide:bis-acrylamide) PAGE in 1⨉ TBE at 80V for 75 min. Gels were imaged on a Typhoon RGB scanner (Cytiva) and signal intensities were quantitated by ImageJ. Background signals (the respective no protein samples) for both the reversal and unwinding products were subtracted from the signal intensities of the reactions with protein.

### AlphaFold modeling

Structural models of the FBH1 and RAD54L interaction were generated using the AlphaFold-Server pipeline. FASTA inputs of human FBH1 (UniProt Q8NFZ0, isoform 1) and human RAD54L (UniProt Q92698) were submitted to the AlphaFold Server. The AlphaFold 3 output models were ranked according to per-residue and interface confidence scores. Model 0 was selected and used for further analyses in UCSF ChimeraX. Using the Matchmaker command in ChimeraX, model 0 was superimposed on the cryo-EM structure of SCFFBH1 at a 3-way DNA replication fork obtained from the Protein Data Bank (PDB ID: 9XZJ, https://doi.org/10.2210/pdb9XZJ/pdb) 11. The contacts command was used to identify residues of RAD54L and FBH1 within 3.5 Å.

## DATA AVAILABILITY

The source data underlying all figures are provided within this paper and in the supplement.

## SUPPLEMENTARY MATERIAL

Supplementary data are available online.

## AUTHOR CONTRIBUTIONS

MEU: Investigation, methodology, validation, formal analysis, writing of the manuscript. CNJ: Critical reagents. JG: Investigation, validation. AVM: Conceptualization, funding acquisition. CW: Conceptualization, formal analysis, writing of the manuscript, funding acquisition.

