## Supporting Information for "FBH1 and RAD54L directly interact and cooperate to drive replication fork reversal"

**Supplemental Figure S1**
**Supplemental Figure S2**
**Supplemental Figure S3**
**Supplemental Figure S4**

**Supplemental Table S1**
**Supplemental Table S2**
**Supplemental Table S3**
**Supplemental Table S4**
**Supplemental Table S5**

### Supplemental Figures

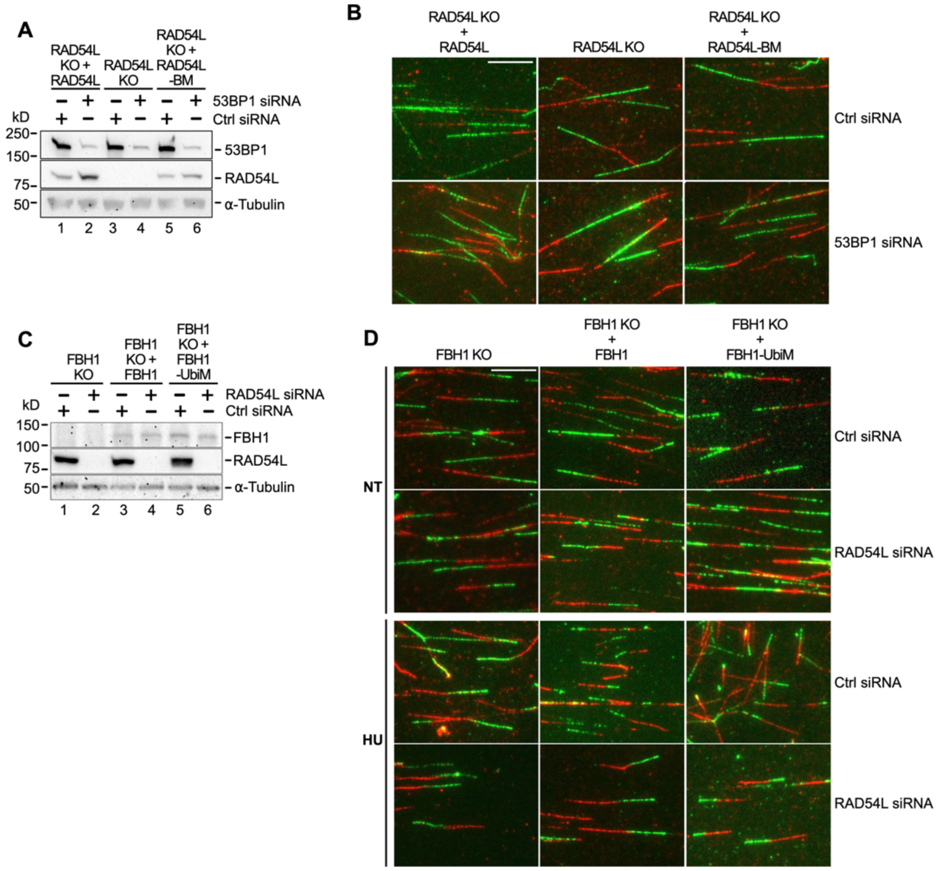


**Figure S1. Related to Figure 1. RAD54L branch migration (BM) is essential in the FBH1 pathway but dispensable for fork reversal by the FBH1 E3 ligase mutant.** (**A**) Representative

Western blots to show extent of 53BP1 knockdown in HeLa RAD54L KO cells and RAD54L KO cells expressing wild type RAD54L or RAD54L-4A/S49E (here: RAD54L-BM) (Fig. 1B, Fig. S1B). Loading control: α-Tubulin. (**B**) Representative micrographs of DNA fibers in HeLa RAD54L KO cells and RAD54L KO cells expressing wild type RAD54L or RAD54L-BM [1] transfected with Ctrl or 53BP1 siRNA (Fig. 1B). (**C**) Representative Western blots to show extent of RAD54L knockdown in U2OS FBH1 KO cells and FBH1 KO cells expressing wild-type FBH1 or FBH1-F266A/P267A (Fig. 1D, Fig. S1D). Loading control: α-Tubulin. (**D**) Representative micrographs of DNA fibers in untreated and HU-treated FBH1 KO cells and FBH1 KO cells expressing wild-type FBH1 or FBH1-F266A/P267A (here: FBH1-UbiM) and transfected with Ctrl or RAD54L siRNA (Fig. 1D). Scale bars: 10 μm.

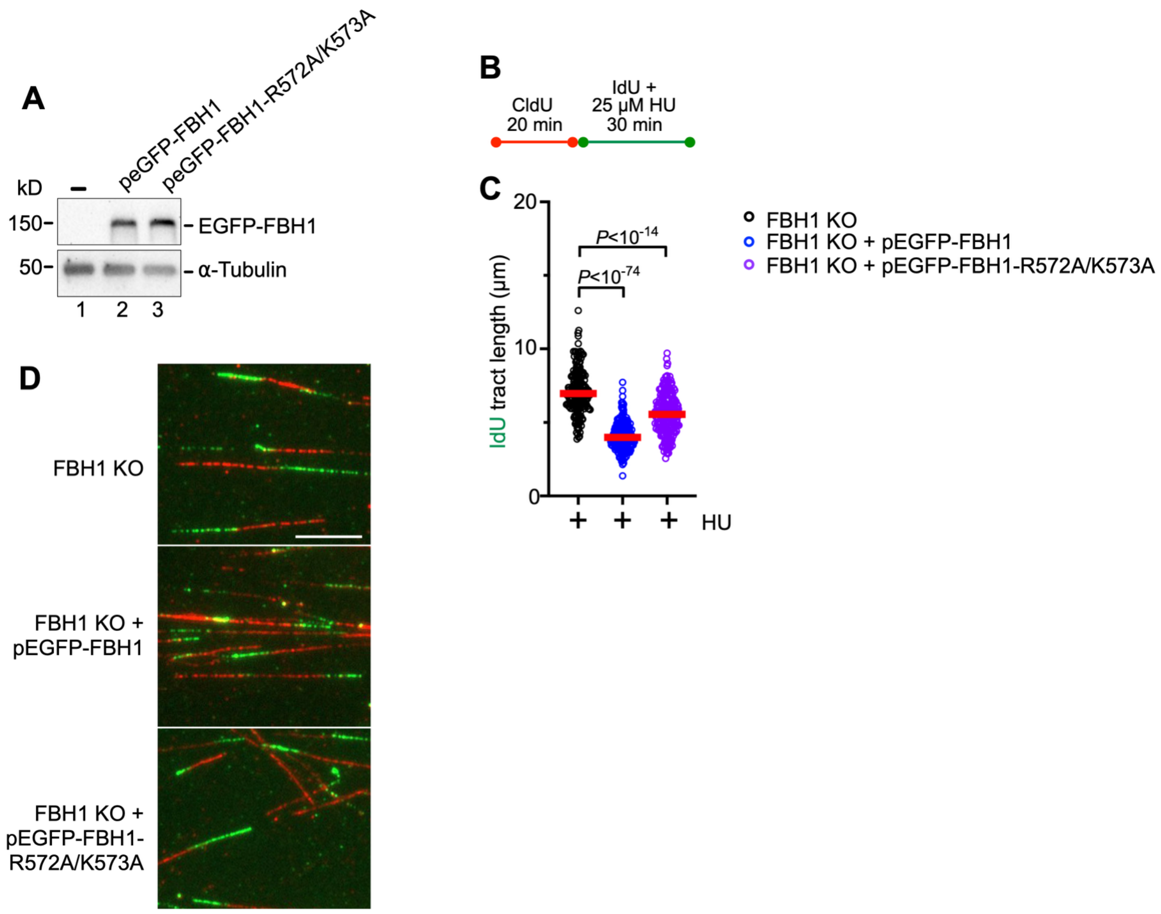


**Figure S2. Related to Figure 3. FBH1 predominantly associates with RAD54L-Lobe 1 in cells, and their association at stalled replication forks is dependent on the identified [2] FBH1 fork junction binding motif.** (**A**) Western blots to show expression of transiently expressed EGFP-FBH1 and EGFP-FBH1-R572A/K573A in RAD54L KO+RAD54L-HA cells. Loading control: α-Tubulin. (**B**) Schematic of the DNA fiber assay protocol used in (C). (**C**) Dot plots with medians (red) of IdU tract lengths in HU-treated U2OS FBH1 KO cells and FBH1 KO cells expressing wild-type FBH1 or mutant FBH1-R572A/K573A (n=2; 206-239 fiber tracts analyzed; Kruskal-Wallis test followed by Dunn's multiple comparisons test). (**D**) Representative micrographs of DNA fibers in HU-treated FBH1 KO cells and FBH1 KO cells expressing wild-type FBH1 or FBH1- R572A/K573A (Fig. S3C). Scale bar: 10 μm.

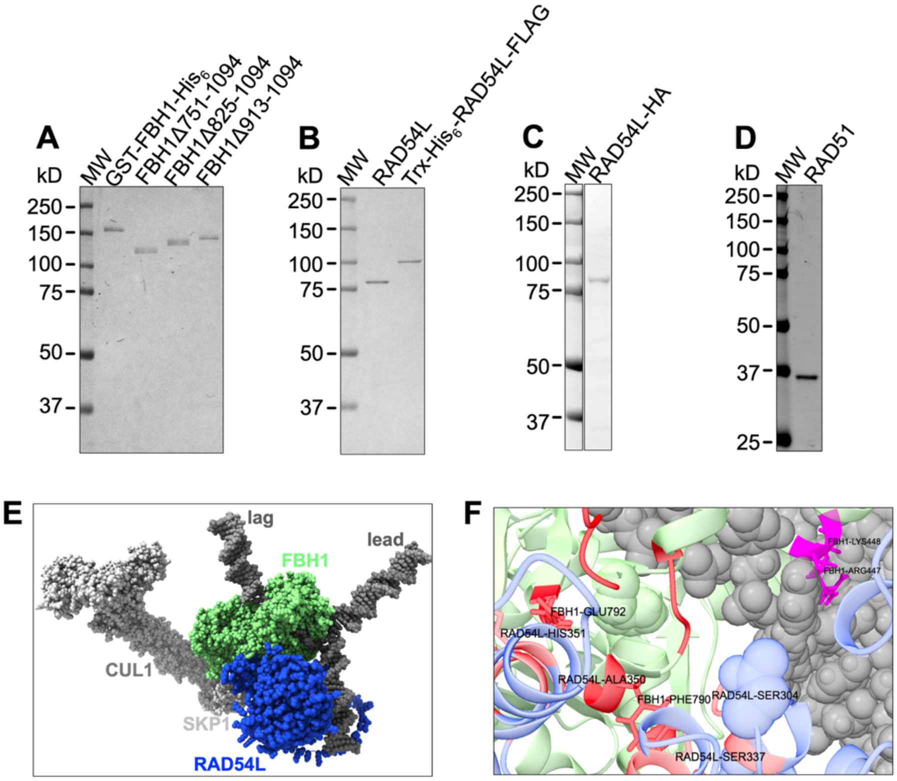


**Figure S3. Related to Figure 4. RAD54L and FBH1 directly interact.** (**A**) SDS-PAGE to show purified GST-FBH1-$His_6$, GST-FBH1Δ751–1049, GST- FBH1Δ825–1049, and GST- FBH1Δ913–1049. (**B**) SDS-PAGE to show purified RAD54L and Trx-$His_6$-RAD54L-FLAG. (**C**) SDS-PAGE to show purified RAD54L-HA. (**D**) SDS-PAGE to show purified RAD51. (**E**) AlphaFold 3 structural prediction of complex formation between RAD54L and FBH1 at a 3-way DNA model replication fork. An AlphaFold 3 model of the interaction between RAD54L and FBH1 was superimposed on the cryo-EM structure of $SCF^{FBH1}$ at a 3-way DNA replication fork (PDB ID: 9XZJ, https://doi.org/10.2210/pdb9XZJ/pdb [2]). (**F**) Zoomed-in view of the AlphaFold 3 model described in (E). Green: FBH1; Blue: RAD54L; Gray: DNA; Red: predicted residues of contact for RAD54L and FBH1 within 3.5 Å; Magenta: R447/K448 (in FBH1 isoform 4) identified as the FBH1 RF junction binding motif [2]; Black labels: residues in RAD54L and FBH1 with predicted contacts in UCSF ChimeraX (FBH1 residues are based on isoform 4).

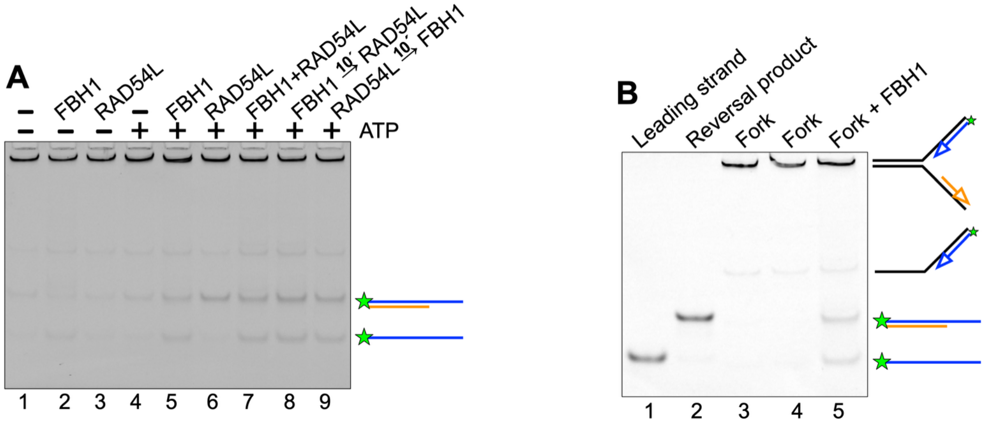


**Figure S4. Related to Figure 5. Replication fork reversal is more efficient with RAD54L and FBH1.** (**A**) Representative gel image of the *in vitro* fork reversal assays with FBH1 and RAD54L used for quantification. (**B**) Gel image of migration markers (lanes 1-4) and fork reversal products by FBH1 (lane 5). The faint band corresponding to one arm of the fork (lanes 3-5) likely is the product of incomplete annealing during substrate preparation.

### Supplemental Tables

**Supplemental Table S1.** Predicted contacts between FBH1 and RAD54L.

| FBH1 Residue (isoform 4) | FBH1 Residue (isoform 1) | RAD54L Residue | Number of Predicted Contacts |
|---|---|---|---|
| E792 | E917 | H351 | 13 |
| I785 | I910 | E305 | 7 |
| F790 | F915 | S304 | 5 |
| F790 | F915 | Y309 | 4 |
| F790 | F915 | H340 | 3 |
| D789 | D914 | H340 | 3 |
| F790 | F915 | F336 | 2 |
| E792 | E917 | K354 | 2 |
| F790 | F915 | S337 | 1 |
| I785 | I910 | S304 | 1 |
| E792 | E917 | A350 | 1 |
| E786 | E911 | Q310 | 1 |
| D789 | D914 | A350 | 1 |
| D787 | D912 | Y309 | 1 |

**Supplemental Table S2.** siRNAs used in this study.

| Name | Target Sequence (listed 5´- 3´) | References |
|---|---|---|
| non-depleting control (Ctrl) | GATTCGAACGTGTCACGTCAA | [1] |
| RAD54L | AAGCATTTATTCGAAGCATTT | [1, 3] |
| RAD51 | GAGCTTGACAAACTACTTC | [3] |

**Supplemental Table S3.** PCR primers used in this study.

| Primer | Sequence (listed 5´- 3´) | Product length (bp) |
|---|---|---|
| Amplifying FBH1-HA to clone into pENTR1A - Fwd | ATCGGGTACCATGAGCTACGAGG | 3419 |
| Amplifying FBH1-HA to clone into pENTR1A - Rev | CGATGTCGACTCAATGGTGGACG | |
| Amplifying FBH1 to clone into peGFP-C1- Fwd | ATCGGTCGACTCAGCTACGAGGTGACTTCAGGCTGCC | 3282 |
| Amplifying FBH1 to clone into peGFP-C1- Rev | CGATGGTACCTCAGAAGACGAGGAAGAGCAGGGCCTCA | |
| Amplifying FBH1 to clone into pGEX - Fwd | ATCGGTCGACTCAGCTACGAGGTGACTTCAGGCTGCC | 3282 |
| Amplifying FBH1 to clone into pGEX - Rev | GAGGCCCTGCTCTTCCTCGTCTTCGGAAGTGGACATCACCATC ACCATCACTGAGCGGCCGCATCG | |
| Primers to clone GST-FBH1-His6 into pENTR1A - Fwd | ATCGGGTACCATGTCCCCTATACTAGGTTATTGGAAAATTAAGG GCCTTGTGCAACCC | 4016 |
| Primers to clone GST-FBH1-His6 into pENTR1A - Rev | CGATGCGGCCGCTCAGTGATGGTGATGGT | |

**Supplemental Table S4.** Oligonucleotides used in this study for site-directed-mutagenesis.

| Template | Oligonucleotide | Target Sequence (listed 5´- 3´) |
|---|---|---|
| peGFP-C1-FBH1 | R572A/K573A forward | gcggcgTACCAGTCAAAGAAGAAGTTG |
| | R572A/K573A reverse | CCTATGTGCCCGTAG |
| pENTR1A-GST-FBH1-His6 | Q751toSTOP forward | CTATCTCACGtgaAGTTTTCGGTTTG |
| | Q751toSTOP reverse | AAGACGTGGGTGTGG |
| pENTR1A-GST-FBH1-His6 | G825toSTOP forward | TCTTCAGCCAtgaGAAGAACGGAGG |
| | G825toSTOP reverse | AGGATCCAAATATCAATGATTC |
| pENTR1A-GST-FBH1-His6 | L913toSTOP forward | GGAGTTTGACtgaGTGCATGTTTTGG |
| | L913toSTOP reverse | AGGCCTTTGGCTTTG |

[1]Mutated bases in lower case characters.

**Supplemental Table S5.** Oligonucleotides used in this study for the *in vitro* reversal assay, adapted from [4].

| Oligonucleotide | Sequence | Details |
|---|---|---|
| #71 (94mer) | CTT TAG CTG CAT ATT TAC AAC ATG TTG ACC TAC AGC ACC AGA TTC AGC AAT TAA GCT CTA AGC CAT CCG CAA AAA TGA CCT CTT ATC AAA AGG A | Parental DNA |
| #117 (94mer) | T CCT TTT GAT AAG AGG TCA TTT TTG CGG ATG GCT TAG AGC TTA ATT GCT GAA TCT GGT GCT GTA GGT CAA CAT GTT GTA AAT ATG CAG CTA AAG | Parental DNA |
| #2 (61mer) | TCC TTT TGA TAA GAG GTC ATT TTT GCG GAT GGC TTA GAG CTT AAT TGC TGA ATC TGG TGC TGT | Corresponding to a 2bp leading strand gap at the fork |
| #1 (55mer) | A GAT TCA GCA ATT AAG CTC TAA GCC ATC CGC AAA AAT GAC CTC TTA TCA AAA GGA | Corresponding to an 8bp lagging strand gap at the fork |
